# Boosting the MS1-only proteomics with machine learning allows 2000 protein identifications in 5-minute proteome analysis

**DOI:** 10.1101/2020.10.29.359075

**Authors:** Mark V. Ivanov, Julia A. Bubis, Vladimir Gorshkov, Daniil A. Abdrakhimov, Frank Kjeldsen, Mikhail V. Gorshkov

## Abstract

Proteome-wide analyses most often rely on tandem mass spectrometry imposing considerable instrumental time consumption that is one of the main obstacles in a broader acceptance of proteomics in biomedical and clinical research. Recently, we presented a fast proteomic method termed DirectMS1 based on MS1-only mass spectra acquisition and data processing. The method allowed significant squeezing of the proteome-wide analysis to a few minute time frame at the depth of quantitative proteome coverage of 1000 proteins at 1% FDR. In this work, to further increase the capabilities of the DirectMS1 method, we explored the opportunities presented by the recent progress in the machine learning area and applied the LightGBM tree-based learning algorithm into the scoring of peptide-feature matches when processing MS1 spectra. Further, we integrated the peptide feature identification algorithm of DirectMS1 with the recently introduced peptide retention time prediction utility, DeepLC. Additional approaches to improve performance of the DirectMS1 method are discussed and demonstrated, such as FAIMS coupled to the Orbitrap mass analyzer. As a result of all improvements to DirectMS1, we succeeded in identifying more than 2000 proteins at 1% FDR from the HeLa cell line in a 5 minute LC-MS1 analysis.

## INTRODUCTION

High throughput and sensitive analytical approaches enabling proteome-wide measurements of large cohorts of samples will open the way for application of mass spectrometry-based proteomics in clinical trials, population proteomics, as well as in emerging areas of drug-to-proteome interactions and metaproteome characterization.^1,2^ Moreover, the needed throughput and depth of proteome coverage in these studies has to be accompanied with protein quantitation consistent across the sample cohorts, which is necessary for these approaches being useful in personalized medicine studies.^3,4^ A recent example on using the large cohort of COVID-19 patients clearly demonstrated the need for ultra-high-throughput proteomics to generate hypotheses about therapeutic targets and aid classification or diagnostic decision making in clinical environments.^5^

The typical MS/MS-based proteomic methods are extremely instrument time consuming, which is partially overcome by expensive multiplexing using isotopic labeling strategies, such as tandem mass tag (TMT).^6^ Nevertheless, a number of recent studies employing the bottom-up proteomic approaches become increasingly focused on increasing the throughput of the proteome-wide analysis. These studies include all steps of the proteome characterization workflow, such as increasing the speed of sample preparation prior to MS-based analysis^7–11^, squeezing data acquisition time to a few minute range^12–14^ and increasing the throughput of data processing^15–20^.

One of the most time-consuming parts of the analysis is MS/MS data acquisition, which requires using long HPLC gradient times to have enough room for sequential isolation of as many as possible precursor ions from each MS1 spectrum for subsequent fragmentation. A number of methods and approaches to MS/MS-free proteome analysis have been proposed and widely explored starting from the early Accurate Mass and Tag (Time) method^21–25^ to a recent truly (e.g., without employing tandem mass spectrometry at any step in the workflow) MS/MS-free realizations^26,27^. These methods rely heavily on the accuracies of both peptide *m/z* measurements and retention time (RT) predictions. The latter is especially important for the MS/MS-free strategy as only retention times contain sequence-specific information in a (*m/z*, RT) space.^28,29^ With advances in development of machine learning algorithms a variety of highly accurate RT prediction models become increasingly available.^30–33^ The latter work is particularly interesting as it shows that DeepLC model’s performance is comparable with the other deep learning-based alternatives, yet, it provides better generalization between different chromatography setups. This feature is particularly useful for MS/MS-free proteomic approaches such as DirectMS1, when the number of peptides available for RT prediction model training is typically small. Therefore, the use of a pretrained DeepLC model for rapid adoption to a dataset obtained for particular separation conditions becomes advantageous.

Previously, we described the DirectMS1 method, in which proteolytic peptide mixture analysis is performed in MS/MS-free mode of acquisition using high resolution mass spectrometry.^27^ Because the method does not employ isolation and fragmentation steps, the time for peptide separation can be reduced significantly to few minutes and the number of MS1 spectra available for processing will only be limited by the acquisition rate of the mass analyzer operating at high mass resolution. For the first time, DirectMS1 demonstrated the capability to identify up to 1000 proteins from a HeLa cell line using only a 5-minute HPLC gradient. Moreover, the average sequence coverage for each identified protein in this method exceeded the one of a standard MS/MS-based approach (even when long gradient is used) by almost an order of magnitude, thus, significantly improving the quantitation. On a proteome-wide scale this kind of analysis efficiency was not considered feasible before because of the whole proteome sample complexity, a lack of sequence specific information in the measured *m/z* values of peptide ions, and the low accuracy of existing phenomenological retention time prediction models. However, advances in high resolution mass spectrometry technology and machine learningbased RT prediction models implemented in DirectMS1’s search engine, allowed the method to exceed the proteome coverage rate per analysis time of a long gradient MS-MS-based proteomics. Yet, the method’s efficiency does not reflect the real information content of the MS1 dataset, in which all eluted and ionizable peptides of the analyzed proteolytic mixture are potentially present in a full dynamic range accessible by current state-of-the-art high resolution mass analyzers.

In this work, we integrated two novel machine learning algorithms into the data processing workflow of DirectMS1 to further improve its efficiency. These algorithms include DeepLC as a retention time prediction model used in the method’s peptide identification and the gradient boosted machine, LightGBM^34^, for scoring peptide-feature matches. Also, the method was upgraded and tested for processing MS1-only data obtained using high resolution mass spectrometry and the high-field asymmetric waveform ion mobility, FAIMS^35^, which was found increasingly applicable in proteomic research^36,37^ as it provides additional separation dimension for peptides at the front end of a mass spectrometer.

## EXPERIMENTAL SECTION

### Samples

All experiments were performed using Thermo Scientific Pierce™ HeLa Protein Digest Standard (P/N 88328) derived from HeLa S3 cell line.

### Data acquisition

LC-MS analysis in this work was performed using Orbitrap Fusion Lumos mass spectrometer (Thermo Scientific, San Jose, CA, USA) coupled with UltiMate 3000 LC system (Thermo Fisher Scientific, Germering, Germany) and equipped with *FAIMS Pro* interface. Short LC gradient method was applied as reported previously^27^. Trap column μ-Precolumn C18 PepMap100 (5 μm, 300 μm, i.d. 5 mm, 100 Å) (Thermo Fisher Scientific, USA) and self-packed analytical column (Inertsil 3 μm, 75 μm i.d., 15 cm length) were employed for separation. Mobile phases were as follows: (A) 0.1 % formic acid (FA) in water; (B) 80 % acetonitrile (ACN), 0.1 % FA in water. Loading solvent was 0.05 % trifluoroacetic acid (TFA) in water. The gradient was from 5 % to 35 % phase B in 4.8 min at 1.5 μL/min. Total method time including column washing and equilibration was 7.3 min. Field asymmetric ion mobility spectrometry (FAIMS) separations were performed with the following settings: inner and outer electrode temperatures were 100 °C; FAIMS carrier gas flow was 4.7 L/min; asymmetric waveform dispersion voltage (DV) was −5000 V; entrance plate voltage was 250 V. Compensation voltages (CV) −50 V, −65 V, and −80 V were used in a stepwise mode during LC-MS analysis. Mass spectrometry measurements were performed in MS1-only mode of acquisition. Full MS scans were acquired in a range from *m/z* 375 to 1500 at a resolution of 120 000 at *m/z* 200 with Automatic Gain Control (AGC) target of 4·10^5^, 1 microscan and 50 ms maximum injection time. Two hundred ng of HeLa digest was loaded on column.

Experimental data obtained previously for 500 ng HeLa injection in both MS1-only MS/MS using Q-Exactive HF mass spectrometer (Thermo Fisher Scientific, San Jose, CA, USA) were taken from PRIDE repository^38^ (repository identifier PXD015669) and used for testing upgraded DirectMS1 software. Supplementary Table S1 provides a list and the annotations for all experimental data used in this work.

### Data Analysis

Raw files were converted into mzML format using *msConvert* from ProteoWizard (v. 3.0.20066).^39^ Peptide features were detected using either Dinosaur^40^ (v. 1.2.0) or in-house developed Biosaur (freely available at https://github.com/abdrakhimov1/Biosaur) (v. 1.0.3) software for data obtained without and with FAIMS, respectively. Proteins were identified by using *ms1searchpy* (v. 2.0.3), which is the search engine of DirectMS1. This software is open-source and freely available at https://github.com/markmipt/ms1searchpy under Apache 2.0 license. Parameters for the search were as follows: minimum 3 scans for detected peptide isotopic cluster; minimum one visible C13 isotope; charges from 1+ to 6+, no missed cleavage sites, and 8 ppm initial mass accuracy. All searches were performed against Swiss-Prot human concatenated database containing 20247 protein sequences and its decoys unless otherwise stated. Results were filtered to 1% protein level false discovery rate (FDR) using target-decoy approach^41^ with its “picked” modification^42^ and “+1” correction^43^.

### Data Availability

The datasets generated and analyzed during the current study have been deposited to the ProteomeXchange Consortium via the PRIDE partner repository with the dataset identifier PXD022094.

## RESULTS AND DISCUSSION

### The workflow

Figure 1 shows details of DirectMS1 method. The method starts with acquiring high resolution peptide ion mass spectra. A mass spectrometer operates in MS1-only mode and simply collects spectra for eluting peptides at the speed determined by the AGC and mass resolution settings. The total number of MS1 spectra acquired during 5-min LC gradient at the mass resolution of 120,000 at *m/z* 200 and AGC of 3*10^6^ ranges from 1,000 to 1,500 depending on the mass spectrometer model.

**Figure 1.**
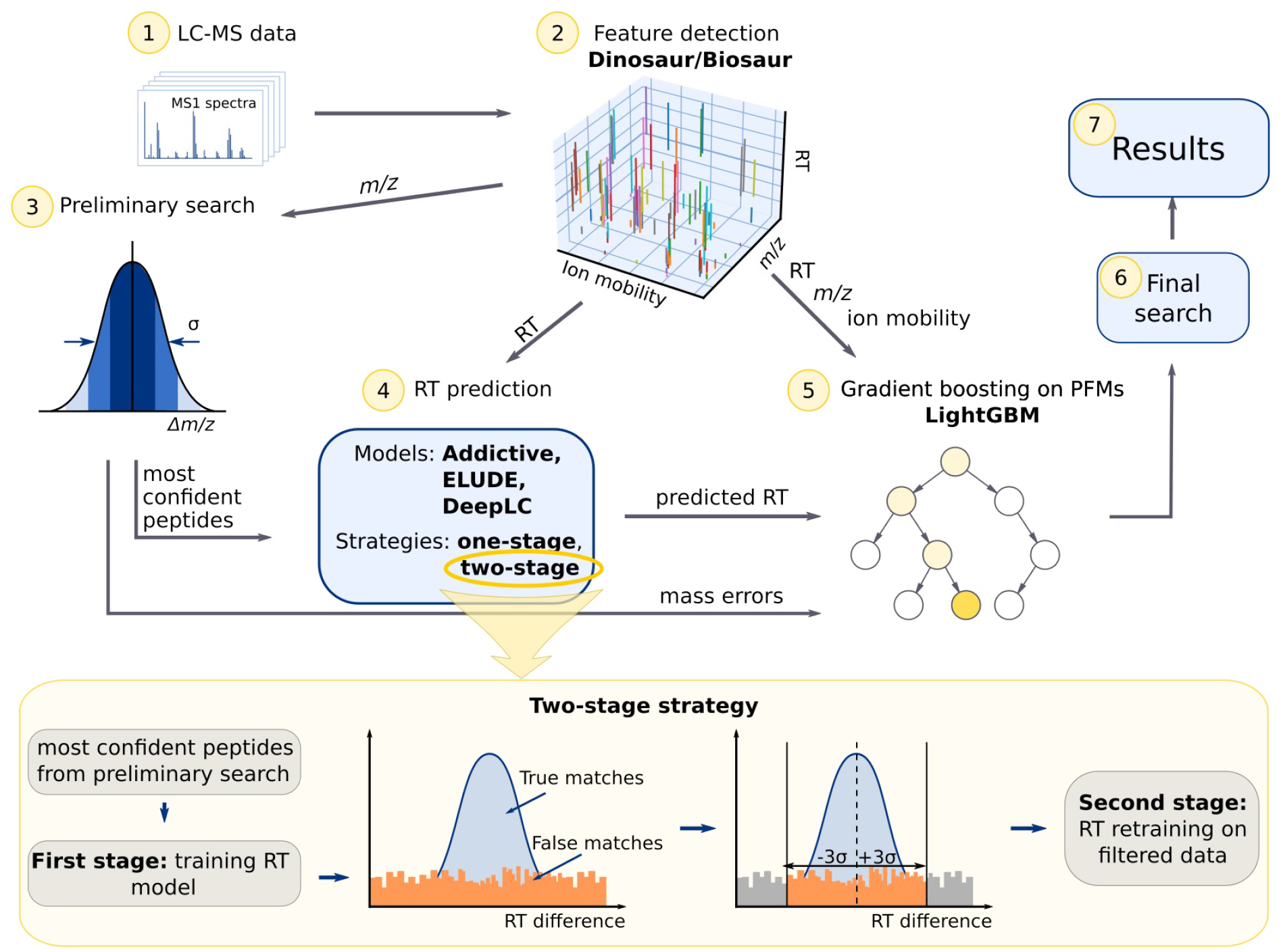
DirectMS1 workflow.

Second step is peptide features detection using either Dinosaur or Biosaur utilities. The latter was developed in-house recently and allows handling ion mobility data, such as FAIMS. The third step is preliminary searchperformed using only the information about measured peptide *m/z* values. High confident peptide feature matches (PFMs), the ones belonging to the top scored proteins identified in the preliminary search and having the smallest mass error, are selected for RT prediction models calibration in the next step. By default, the first 2500 PFMs with the highest confidence are used, many of them being false matches (see discussion below).

Training of peptide RT prediction models integrated into the search workflow is one of the crucial steps in DirectMS1. Originally it was having an option to use one of the two RT models. The first one was relying on the so-called “additive” retention time prediction model, in which a peptide retention time is a sum of individual retention coefficients of residues constituting the peptide sequence.^44^ These coefficients can be either predefined for the type of used chromatography, or calculated on-the-fly for specific data and separation conditions. The second model was based on the ELUDE^30^ RT prediction software and in general allows significantly more accurate prediction of peptide retention times compared with the additive model, although for the price of longer computation time. In this work we upgraded DirectMS1 search module by integrating it with the recently introduced deep learning RT prediction model - DeepLC^33^. It performs reportedly as well, or better, as the other state-of-the-art retention time prediction models employing machine learning algorithms. Indeed, DeepLC model allowed significant improvement in the accuracy of retention time prediction for identifying peptides in DirectMS1 compared with the previously used additive and ELUDE models that resulted in dramatic increase in the number of identified proteins as shown in Figure 2a and b. Furthermore, a two-stage calibration strategy was implemented, which allowed to further decrease RT prediction error and increase the number of identifications as detailed below in the *Retention time prediction models* section.

**Figure 2.**
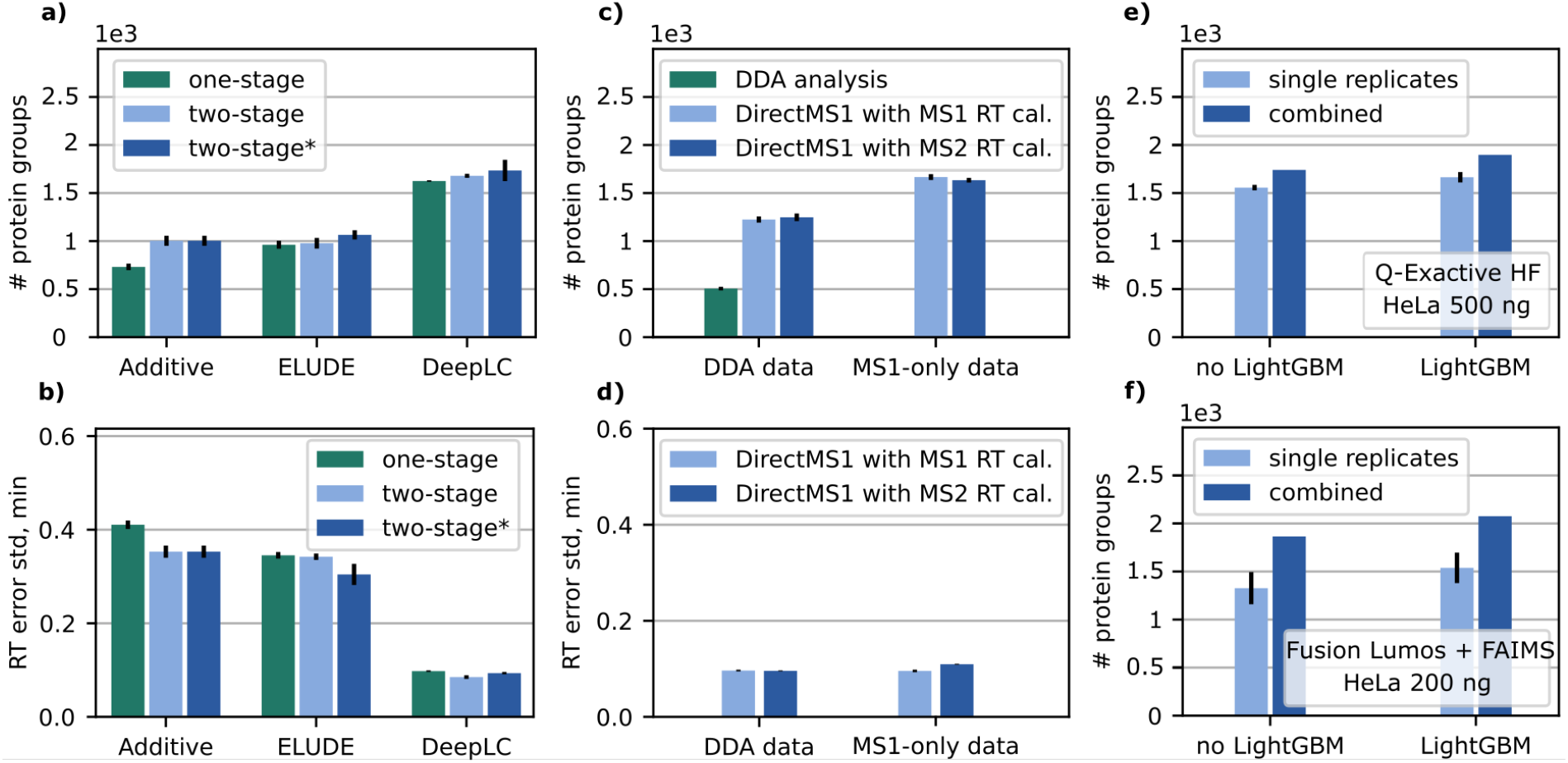
*Left column*. Analysis of HeLa cell line runs using DirectMS1: **(a)** Number of protein groups identified at 1% FDR for different RT training strategies.”One-stage” denotes standard RT training, “two-stage” -- RT training using two-stage strategy with the same prediction model for both stages, and “two-stage*” -- RT training using additive and DeepLC models at first and second stages of training, respectively; (**b)** Retention time prediction accuracy determined as a standard deviation in the distribution of peptide matches of identified proteins by the difference between experimental and predicted RT for different RT training strategies. *Middle column*. performance of DirectMS1 with different initial training sets for RT prediction: (**c)** Identified protein groups at 1% FDR; and (**d)** Standard deviation of difference between experimental and predicted retention time for reliable peptide matches. The results are shown for DirectMS1 analysis of HeLa cell line using LC-MS1-only and DDA data with excluded MS/MS spectra. Green color is DDA MS/MS analysis results. *Right column*. Number of identified protein groups at 1% FDR for DirectMS1 with gradient boosting model, LightGBM, turned on/off: (**e)** Q-Exactive HF; and (**f)** Fusion Lumos with FAIMS setup. Results are shown in average for 3 replicates, as well as for all replicates combined into single identification results.

At the fifth step of the workflow, a gradient boosting algorithm is applied for estimating the PFMs quality. The final search uses PFMs quality information for protein score calculations and prepares protein identification list at the specified FDR level, typically 1% (see discussion below).

### Retention time prediction models

Retention time prediction procedure in DirectMS1 has an important distinction compared with the typically used in conventional MS/MS-based searches. The latter usually uses a list of high-confident peptide-spectrum matches (PSM) (typically filtered to 1% FDR level) for RT model training, retraining, or calibration. On the contrary, identifications in DirectMS1 have a high rate of false positives among PFMs and it is not possible to obtain a subset of high-confident peptides of the size reasonable for RT training. DirectMS1 begins with a preliminary search as shown in Fig.1 as the second step of the workflow using only *m/z* values of detected PFMs. Then, it forms a preliminary list of identified proteins at 1% FDR. Each of these proteins contains a set of PFMs, with some of them being wrong, and these PFMs have different levels of deviations of experimentally determined *m/z* from the theoretically calculated one. PFMs with the lowest mass deviations, up to 2500 PFMs in case of the cellular proteome analyses, are further taken for training the DeepLC model. Also note that DirectMS1 uses RT prediction models in different ways. While the additive model is trained from scratch with no use of previously trained models, both ELUDE and DeepLC are the pretrained models re-calibrated for the current data. Importantly, the results of these recalibrations can be affected by the presence of false positives matches in the training set of PFMs. To alleviate this problem, DirectMS1 employs two-stage RT calibration as shown in Figure 1, bottom panel. First, the model is trained using all 2500 peptides from the preliminary training set. Then, the software calculates the differences between predicted and experimental retention times and fits this difference array with the sum of Gaussian and uniform distributions. Here, we assume that true peptide matches should follow the Gaussian distribution, while the false matches follow the uniform one. Using the standard deviation obtained for the distribution of all PFMs by differences between experimental and predicted RTs, peptide matches with RT differences exceeding three standard deviations are excluded from the training set. Then, the model is subjected to final training using the updated PFM set. Figure 2a and b illustrate how different RT prediction models and one- or two-stage training procedures affect the performance of DirectMS1 method. The results show that the additive model benefits the most from the two-stage RT training process, although it still underperforms compared with machine learning based approaches, which are probably less influenced by the presence of false matches in the training set. However, the two-stage RT training where the first stage is performed using the additive model followed by training DeepLC by the set of RT filtered PFMs as described above provides the maximal number of identified proteins. This two-stage training involving two different RT prediction models is denoted as *two-stage\** in Figure 2 and is currently used by default in DirectMS1’s search engine. All further results presented here for demonstration of the efficiency of the method have been obtained using the two-stage* training.

### Peptide set for RT prediction models

DirectMS1 identifies proteins without use of any MS/MS spectra. However, one can expect that having MS/MS-based identifications for RT model training may potentially benefit the DirectMS1 analysis. This training could be done in two ways: (i) all data are acquired in DDA mode and MS1 spectra are then processed by DirectMS1, while MS/MS identifications are used for RT model calibration; and (ii) a separate single DDA run is performed, which is used for RT model training for several subsequent MS1-only experiments. Both approaches have some drawbacks. Separate DDA run require additional experimental time and sample and can also result in RT alignment errors.^45^ On the other hand, analyzing data using standard DDA mode reduces the number of available MS1 spectra, reducing DirectMS1 performance, or requires longer gradient times for the whole proteome characterization undermining the concept of high throughput proteomics. Also, both abovementioned approaches for RT model calibration require a mass spectrometer capable for MS/MS mode, while DirectMS1 method can be used with simpler and more robust MS1-only instrumentation. To further elaborate on the issue we calculated the number of identified protein groups at 1% FDR and RT prediction error obtained using DeepLC model for different approaches as shown in Figure 2c and d. Firstly, DirectMS1 applied to DDA data shows that RT calibration using peptides identified in MS1 analysis is as good as the one based on the peptides identified from MS/MS spectra. Secondly, DirectMS1 applied to MS1-only data shows that transferring RT calibration results between datasets decreases the overall RT prediction accuracy.

### Machine learning analysis of PFMs

Distributions of PFMs’ mass and RT errors are shown in Figure 3 a, b for target and decoy matches. Expectedly, the mass error does not provide a strong differentiation between target and decoy PFMs as shown in Fig 3a. On the other hand, retention time difference between experimental and predicted values present potentially a strong discrimination metric since peptide retention times are sequence-specific (Figure 3b). Note also, that there can be some other peptide properties discriminating targets and decoys. For example, the RT prediction and mass measurement errors can depend on peptide lengths and signal intensities, respectively. Gradient boosting techniques should potentially account for these non-linear effects and use them to increase the discriminative power of the search engine. In DirectMS1’s search engine *ms1searchpy* we employed the LightGBM gradient boosting framework for estimating the PFM quality score. A list of target-decoy discriminative properties used for LightGBM in the current version of *ms1searchpy* is shown in Supplementary Table S2. Hyperparameters for LightGBM are tuned in DirectMS1 on-the-fly during data processing. It is implemented using *Random Search Cross-Validation method* with 25 random hyperparameters testing and 3-fold group based cross-validation. Unique peptide sequences are chosen as groups to prevent overfitting by the presence of the same sequences in both training and test crossvalidation sets. The optimal hyperparameters which provide the maximal average number of PFMs at 25% FDR in the test groups are chosen for final LightGBM model training. Calculated values from gradient boosting are further used in the iteration procedure for protein score calculation. Firstly, 10% of PFMs with best LightGBM scores are picked and protein binomial scores are calculated using only these PFMs. At the next iteration, 20% of PFMs are picked and scores are calculated again. It continues with 10% step up to 100%, and in the end 10 intermediate protein scores become available for each protein at the end. These scores are averaged and used as the final protein score for target-decoy filtering and generating the list with protein identification results.

**Figure 3.**
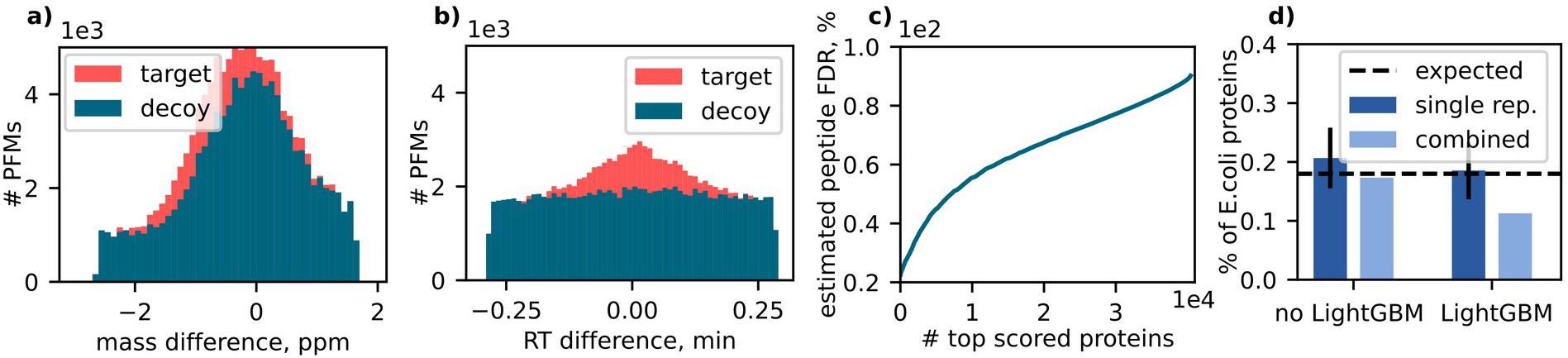
DirectMS1 analysis for 5 min LC-MS data of 200 ng of HeLa lysate using Lumos with FAIMS interface: **(a)** distributions of target and decoy PFMs for mass difference between theoretical and experimental values; **(b)** distributions of target and decoy PFMs for retention time difference between experimental and predicted values; **(c)** estimated peptide-level FDR for a cumulative set of top scored proteins; **(d)** percentage of *E.Coli* proteins in the results at 1% FDR. Here, light blue color is the results for single replicates averaged over 3 technical replicates, and dark blue color is the results for a combination of 3 technical replicates before protein score calculation. Dashed black line is the expected percentage of E. Coli proteins in the results which is a fraction of E. Coli proteins in the search database.

Comparison of DirectMS1 performance with and without the LightGBM framework is shown in Figure 2e. Using this framework adds 7 to 10% of protein identifications. Combining multiple replicates into the single output increases further the number of protein identifications at 1% FDR to nearly 1900 for this particular dataset.

### FAIMS data

FAIMS is an atmospheric pressure ion mobility technique that separates gas-phase ions by the difference in their mobility in high and low electric fields. FAIMS separation is orthogonal^36^ to both LC and MS and provides additional on-line fractionation to further improve dynamic range of the analysis of complex samples. Having this third separation dimension in addition to masses and retention times can be beneficial for DirectMS1 method despite the decrease, albeit marginal, in the average number of MS1 scans per peptide feature when multiple compensation voltages are used. Although the prediction models for FAIMS data are yet to be developed, adding this dimension into the DirectMS1 workflow resulted in the meaningful increase in the number of identified proteins. As shown in Fig.2f, we were able to identify 2100 proteins at 1% FDR in 5-min HeLa proteome analysis after combining 3 technical replicates.

### Validation

False discovery rate at peptide level can be estimated by the following formula:

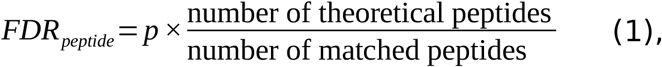

where

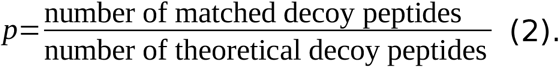

False discovery rate analysis at peptide level is shown in Figure 3c. Strikingly enough, the average FDR for all peptide-feature matches is 90 %. However, the average peptide-level FDR values are 23 % and 32 % for top 100 scored proteins and all proteins at 1% protein FDR level, respectively.

While protein FDR level is controlled by target-decoy approach, the other interesting question is how many proteins from the target database not present in the sample can be found in the results by DirectMS1’s search engine? To perform this control analysis, HeLa data was searched against combined human and *E. coli* Swiss-Prot databases. The number of identified *E. coli* proteins in DirectMS1 analysis is shown in Fig.3d. The expected percentage of *E. coli* proteins is calculated by the formula:

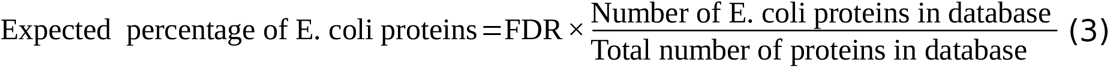

from which we found that 0.18% of *E. coli* proteins at 1% FDR in the identification results as expected (*E. coli* and human databases contain 8862 and 40386 sequences including decoys, respectively). This result shows that DirectMS1 provides an accurate threshold for filtering protein identifications.

## CONCLUSIONS

DirectMS1 for high-throughput MS/MS-free quantitative proteome analysis was upgraded with DeepLC, the novel machine learning-based utility for peptide retention time prediction and the recently introduced gradient boosting framework LightGBM for scoring peptide feature matches in MS1 spectra. Additionally, two-stage and two-model peptide retention time prediction was included into the method’s workflow. Finally, DirectMS1 was tested using a high-resolution mass spectrometer with FAIMS interface that brought the third, peptide structure-dependent dimension into the MS1-only search space. These additions resulted in a two-fold increase in DirectMS1 method’s efficiency which is now capable of identifying as much as 2000 proteins at 1% FDR in a 5-min MS1-only analysis of cellular proteomes. Further improvement in the depth of proteome coverage using this method are expected with improvements in ion mobility data processing and prediction.

## ASSOCIATED CONTENT

### Supporting information

**Supplementary Table S1**. Raw files description with experimental parameters.

**Supplementary Table S2**. A list of target-decoy discriminative properties used for LightGBM.

## ACKNOWLEDGMENTS

The authors thank Prof. Lukas Kall and Dr. Matthew The for helpful discussion and suggestions regarding machine learning analysis of PFMs. Proteomic data processing, statistical analysis, and software development were performed with financial support from Russian Science Foundation, Grant no. 20-14-00229 to M.V.G. Proteomics and mass spectrometry research at University of Southern Denmark SDU are supported by generous grants to the VILLUM Center for Bioanalytical Sciences (VILLUM Foundation grant no. 7292) and PRO-MS: Danish National Mass Spectrometry Platform for Functional Proteomics (grant no. 5072-00007B).

